# Aggressive Periodontitis with Neutropenia Caused by *MMD2* Mutation

**DOI:** 10.1101/827675

**Authors:** Noriyoshi Mizuno, Hiroyuki Morino, Keichiro Mihara, Tomoyuki Iwata, Yoshinori Ohno, Shinji Matsuda, Kazuhisa Ouhara, Mikihito Kajiya, Kyoko Suzuki-Takedachi, Yusuke Sotomaru, Katsuhiro Takeda, Shinya Sasaki, Ai Okanobu, Tetsushi Sakuma, Takashi Yamamoto, Yukiko Matsuda, Ryousuke Ohsawa, Tsuyoshi Fujita, Hideki Shiba, Hideshi Kawakami, Hidemi Kurihara

**Affiliations:** Department of Periodontal Medicine, Graduate School of Biomedical and Health Sciences, Hiroshima University, 1-2-3, Kasumi, Minami-ku, Hiroshima, 734-8553, Japan; Department of Epidemiology, Research Institute for Radiation Biology and Medicine, Hiroshima University, Hiroshima, 1-2-3, Kasumi, Minami-ku, Hiroshima, 734-8553, Japan; Department of Hematology and Oncology, Research Institute for Radiation Biology and Medicine, Hiroshima University, Hiroshima, 1-2-3, Kasumi, Minami-ku, Hiroshima, 734-8553, Japan; Department of Stem Cell Biology, Research Institute for Radiation Biology and Medicine, Hiroshima University, Hiroshima, 1-2-3, Kasumi, Minami-ku, Hiroshima, 734-8553, Japan; Natural Science Center for Basic Research and Development, Hiroshima University, Hiroshima, 1-2-3, Kasumi, Minami-ku, Hiroshima, 734-8553, Japan; Department of Biological Endodontics, Graduate School of Biomedical and Health Sciences, Hiroshima University, 1-2-3, Kasumi, Minami-ku, Hiroshima, 734-8553, Japan; Department of Mathematical and Life Sciences, Graduate School of Science, Hiroshima University, 1-3-1 Kagamiyama, Higashi-Hiroshima 739-8526, Japan

## Abstract

Aggressive periodontitis causes rapid periodontal tissue destruction and is a disease that occurs at a young age and runs in the patient’s family. Here, we revealed a heterozygous A116V missense mutation in the gene encoding monocyte to macrophage differentiation associated 2 (MMD2) protein in a Japanese family with aggressive periodontitis and neutropenia. Analyses of patients’ peripheral blood revealed a low number of neutrophils but abundant quantity of CD34^+^ hematopoietic stem and progenitor cells (HSPCs). Moreover, mutant *Mmd2* mice showed severe alveolar bone loss and neutropenia. In patients and mutant *Mmd2* mice, differentiation of HSPCs into granulocytes was also impeded, and their granulocytes were functionally impaired. Taken together, A116V mutation in *MMD2* gene induced mild neutropenia and slightly limited the immune defense response. Our studies suggested that aggressive periodontitis in association with A116V *MMD2* mutation constitutes a new immune system defect that belongs to the same spectrum of severe congenital neutropenia.

## Introduction

Aggressive periodontitis, formerly called early onset periodontitis or juvenile periodontitis, has an early onset and runs in the patient’s family (Nishimura et al., 1990; Trevilatto et al., 2002; Llorente, 2006). This disease is characterized by the loss of many teeth due to rapid periodontal tissue destruction, with no evident symptoms in other tissues. The prevalence of this disease is 0.1%-0.2% in Caucasians, 0.4%-1.0% in Asians, and 1.0%-3.0% in African Americans (Albandar, 2000). It has been indicated that neutrophil abnormalities lead to the onset of the disease because they allow for bacterial growth, which follows severe periodontal destruction. Especially, abnormalities in neutrophil chemotaxis and superoxide production have been reported as the causes, but the detailed mechanism has not been elucidated (Van Dyke, 1980; Van Dyke, 1985; Shapira, 1991). The current treatment of this disease is mainly symptomatic, but some patients have a poor outcome even after treatment. The purpose of this study was to identify the causative gene of aggressive periodontitis and analyze whether the onset of this disease was associated with neutrophil abnormalities in order to elucidate its pathogenesis.

Here, we reported the identification of a missense mutation in the *monocyte to macrophage differentiation associated 2 (MMD2)* gene, in a Japanese family with autosomal dominant aggressive periodontitis. Furthermore, we provided some insight about the involvement of MMD2 protein in neutrophil differentiation.

## Results

### Characteristic findings in patients

In this study, we focused on a Japanese family with dominantly inherited aggressive periodontitis. The pedigree of this family is shown in Fig. 1A. The proband (III-4) developed gingival swelling and pain in his late teens but was admitted at our hospital at the age of 24. His upper left molar was difficult to preserve and thus it was extracted. Deep periodontal pockets were found in other teeth. The proband’s two brothers (III-2 and III-3) also showed the same symptoms in their late teens. In a further interview, it was noted that proband’s deceased grandfather (I-1), deceased father (II-4), and uncle (II-6) also had aggressive periodontitis, and the deceased I-1 and II-4 individuals used dentures from a young age. Interestingly, the patients were systemically healthy, except they had severe periodontitis with alveolar bone loss. CT images of subjects III-2 and III-4 at the age of 45 and 40, respectively, exhibited a damaged tooth with half to one-third alveolar bone resorption around the tooth root, compared with that of the healthy subject, even though the patients had received professional dental treatment from teen-age to their 40s (Fig. 1B). On the other hand, the examination of oral cavities of II-1, II-5, II-9, and III-5 revealed that they were healthy and not suffering from periodontitis. Data on complete blood count (III-2, 3, and 4) are given in Table S1. Red blood cell and platelet counts were normal; meanwhile, the white blood cell counts were below the lower normal limit. The ratios of neutrophils in the white blood cells were also relatively lower, the numbers were approximately 883-970 /µl. Flow cytometric analysis revealed adequate amounts of CD34^+^ hematopoietic stem and progenitor cells (HSPCs) (III-2; 2.6 ± 1.07 %, III-4; 1.74 ± 0.53 %) in the patients’ bone marrow (Ueda et al., 2001). Even though the numbers of CD34^+^ HSPCs in the bone marrows of the two patients were normal, their differentiation to granulocytes were impeded (Fig. 2A). We observed that apoptotic cells were higher in number in these patients’ setting (data not shown). Interestingly, detailed flow cytometric analysis indicated many CD34^+^ HSPCs in patients’ peripheral blood (CD34^+^ HSPCs/ CD45^+^ cells accounted for 3.3 % in patient III-2, 1.7 % in patient III-4, and 0.08 % in age-matched healthy subject) (Fig. 2B), which is only transiently seen when peripheral blood is mobilized by granulocyte colony stimulating factor (G-CSF) and high-dose of chemotherapy or plerixafor. However, this condition seen in patients’ peripheral blood was persistent rather than transient. Moreover, neutrophil chemotaxis, assessed by stimulation with N-formyl-methionyl-leucyl-phenylalanine (fMLP), was also decreased in the analyzed patients when compared to that of age-matched healthy subjects (Fig. 2C).

**Fig. 1.**
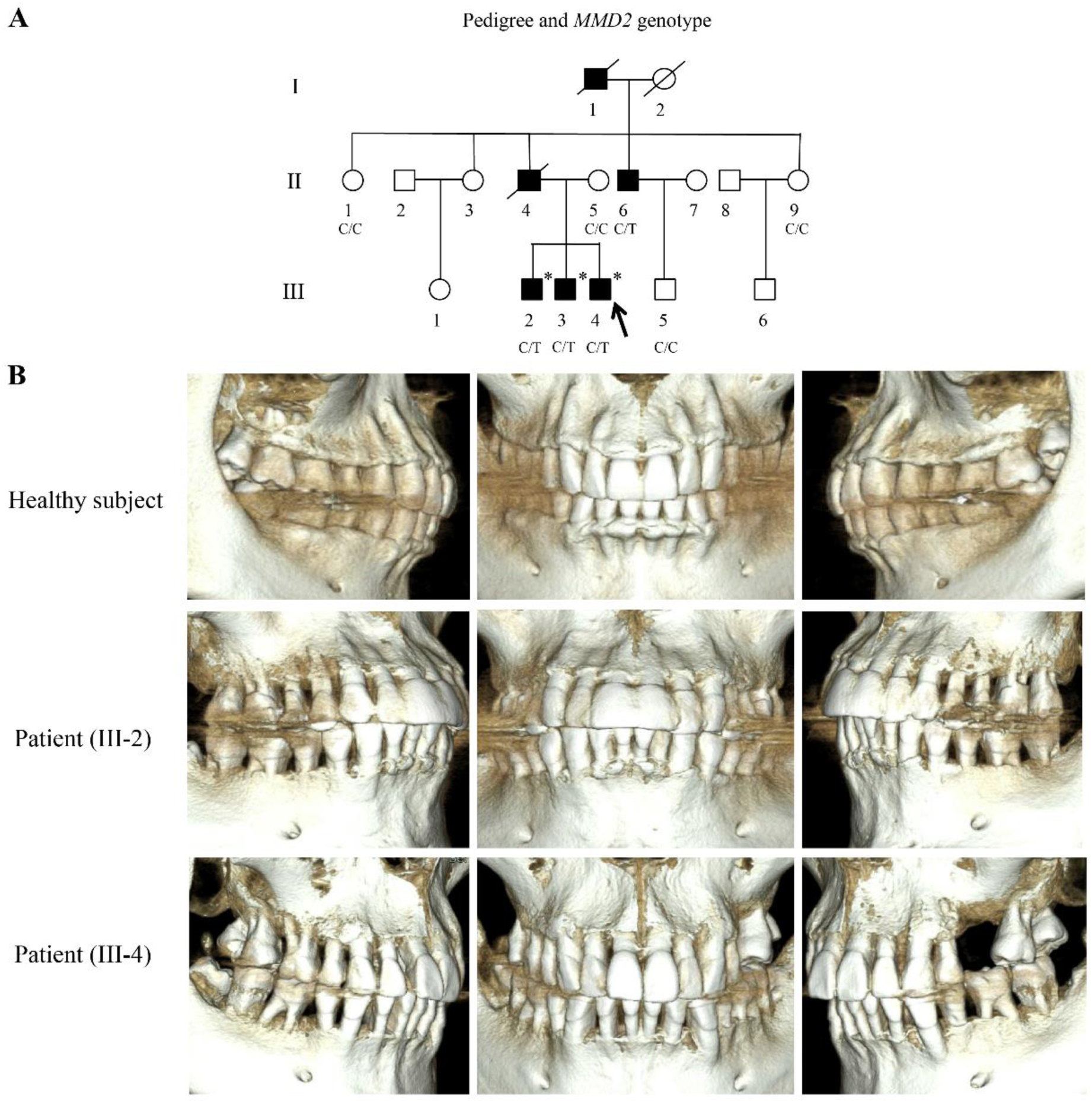
Characteristic findings in patients with *MMD2* mutation. Panel A shows the family tree chart. Arrows indicate the proband. Filled and open symbols represent affected and unaffected individuals, respectively. Genotypes of the variant c.347C>T are shown under the number of samples. Asterisks indicate the patients whose samples were used for exome sequencing. Panel B shows computed tomography images of age-matched healthy subjects (upper), a 45-year-old III-2 patient (middle), and a 40-year-old age III-4 patient (lower).

**Fig. 2.**
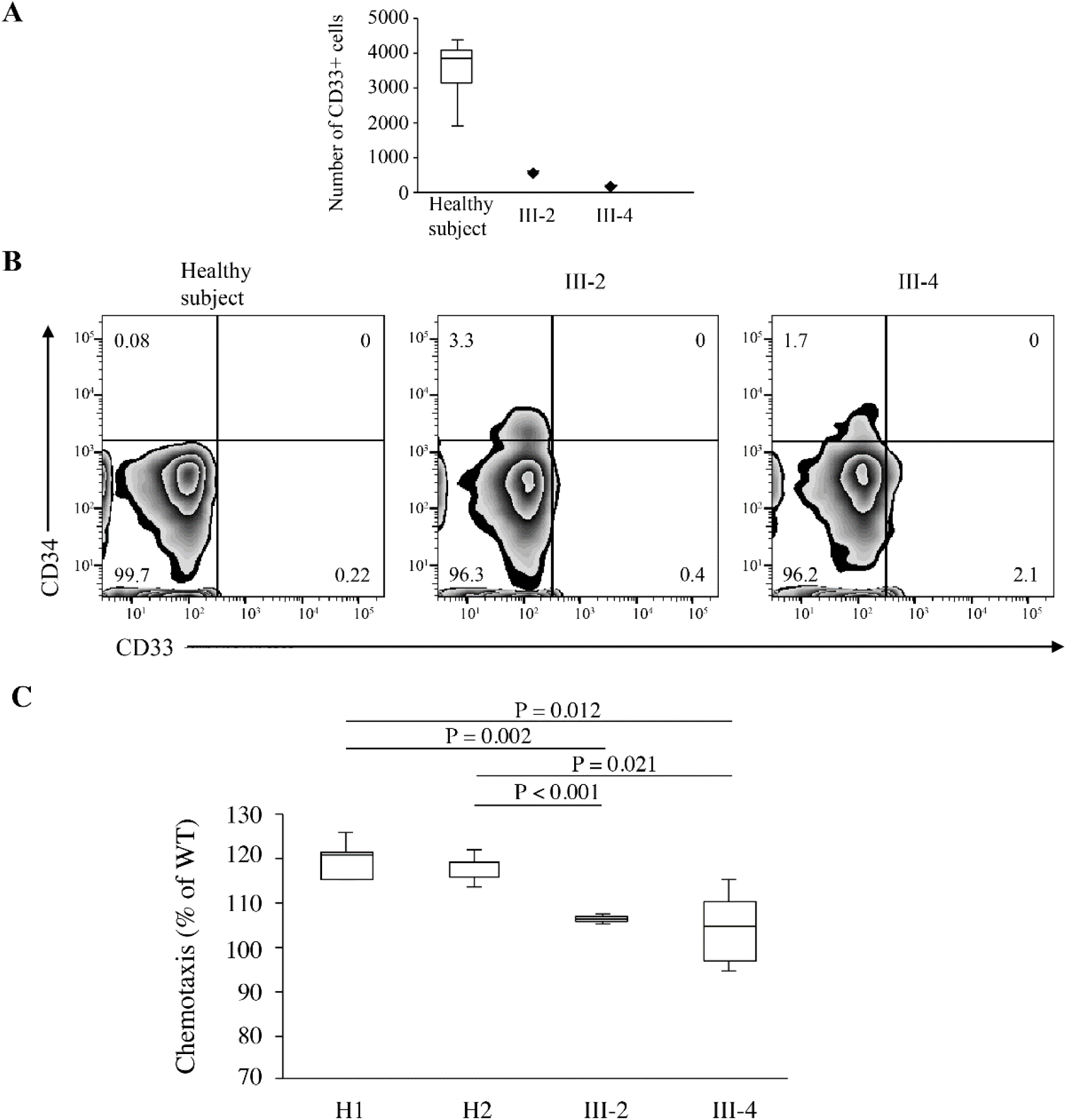
Cellular analysis in patients with *MMD2* mutation. Panel A indicates the induced number of CD33^+^ cells from CD34^+^ HSPCs of the patients III-2 and III-4, which decreased in number compared to that of healthy subjects. T bars indicate standard deviations. Panel B shows flow cytometric analysis of HSPCs from healthy subjects and (III-2 and III-4) patients after gating on CD45^+^ cells and by using anti-CD33 and anti-CD34 antibodies. Panel C shows that neutrophil chemotaxis, induced by fMLP (100 nM, 4 h), in the patients III-2 and III-4 was decreased compared with that of healthy subjects (H1, H2). The results are expressed as the mean ± standard deviation. Those p-values ≤ 0.05 were considered statistically significant by using Student’s t test.

### Identification of the *MMD2* mutation

Our aim was to identify the causative gene mutation in this family. By the linkage analysis, two regions with logarithm of odds (LOD) score of 1.8047 and 1.8058 were obtained in chromosomes 3 and 7, respectively (Fig. 3A and B). As a result of exome sequencing, seven heterozygous variants were identified (Table S2). We performed segregation analysis of the variants in the eight subjects from the family. The variant c.347 C>T, p. A116V in *MMD2* was observed only in affected subjects (II-6, III-2, III-3, and III-4) but not in unaffected subjects (II-1, II-5, II-9, and III-5). Based on the linkage analysis, this *MMD2* variant was present in the high LOD region; therefore, we concluded that *MMD2* variant was most likely associated with periodontitis. This variant was further confirmed by Sanger sequencing (Fig. 3C). The nucleotide and amino acid sequences of the mutated region were completely conserved among vertebrates (Fig. 3D). This variant had a Combined Annotation Dependent Depletion (CADD) score of 28.9 and was pathogenic (>15) (Table S3). This *MMD2* variant was neither found in other Japanese 102 patients with severe periodontitis nor in 275 healthy subjects. Additionally, this mutation has not been detected in the Integrative Japanese Genome Variation Database (database for 7,108 allele number), Human Genetic Variation Database (database for 2,416 allele number), or East Asian Database (database for 19,530 allele number, allele frequency; 0.000), but it was found in 8 alleles in European (database for 128,298, allele frequency; 0.00006235) and in 2 alleles in Latino populations (database for 35,374, allele frequency; 0.00005654) by Genome Aggregation Database. Therefore, this mutation is rare, but it exists.

**Fig. 3.**
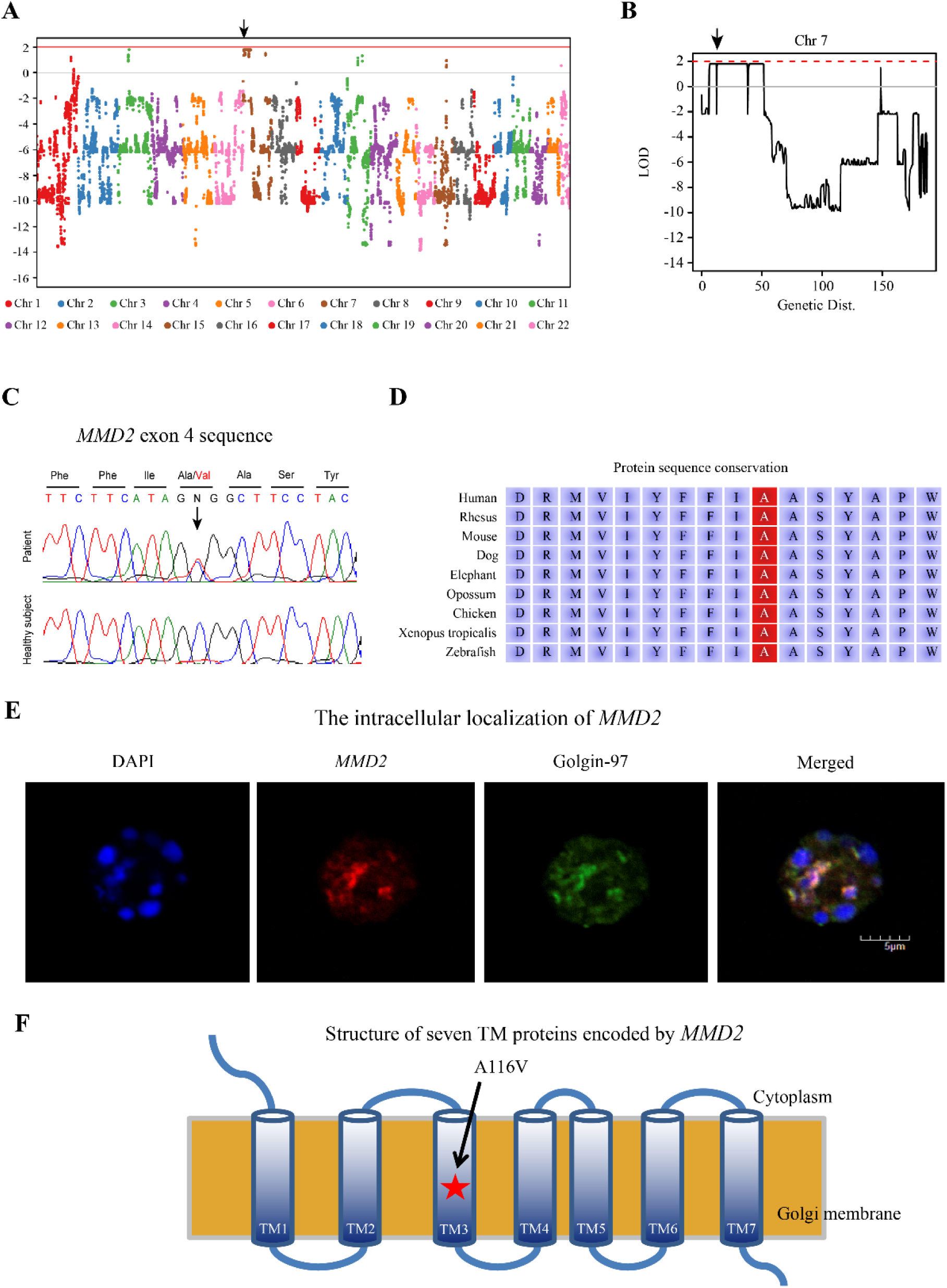
Identification of the *MMD2* mutation. Panels A and B show linkage analysis of the studied family. Arrows indicate the position of *MMD2* gene. Panel C shows Sanger sequencing of *MMD2* gene exon 4 with or without the mutation. Panel D specifies the amino acid sequences that were completely conserved among vertebrates. Panel E revealed the intracellular localization of MMD2 protein in neutrophil-like HL-60 cells. The Golgi was labeled with Golgin-97 (green); meanwhile, nuclei were stained blue with DAPI. Cells were observed under a confocal fluorescence microscope. Panel F shows the structure of seven TM domains encoded by *MMD2*. The star indicates the position of the identified mutation in TM3.

### Cellular localization of MMD2 protein

Immunohistochemical studies were performed to examine the subcellular localization of MMD2 protein. Although human MMD2 protein localization is restricted to the Golgi apparatus when overexpressed (Jin et al., 2012), in our study we observed its presence in the Golgi apparatus even in a steady state, colocalizing with the Golgi marker Golgin-97 (Fig. 3E) in neutrophil-like HL-60 cells. MMD2 is known as an integral membrane protein with its N-terminus facing the cytosol and seven transmembrane (TM) regions (Tang et al., 2005). Specifically, the *MMD2* A116V mutation was localized in TM3 (Fig. 3F).

### Mouse model

The exact function of MMD2 has not been elucidated. Therefore, we created a knock-in mouse model (*Mmd2*^A117V/A117V^ mice) carrying an amino acid substitution in Mmd2, which corresponded to the A116V mutation observed in the *MMD2* human gene. Platinum TALENs were designed to generate a double-stranded break near the targeted nucleotide c.347C>T in exon 4 of *Mmd2* gene (Fig. 4A). The target template for homologous recombination was constructed (Fig. 4B). The 25 bp oligonucleotide contained 4 single nucleotide differences, including the nonsynonymous C>T substitution encoding the mouse A117V mutation and a synonymous change that introduced a PstI site (Fig. 4C). The genome editing of the targeted nucleotide mutations resulted in five random deletions and one indel, as the 5 bp, 7 bp, and 11 bp deletions produced frameshift mutations. We used the mice with 7 bp deletions in *Mmd2* as the knock-out mice (*Mmd2*^-/-^ mice) (Fig. 4D and Fig. 4E).

**Fig. 4.**
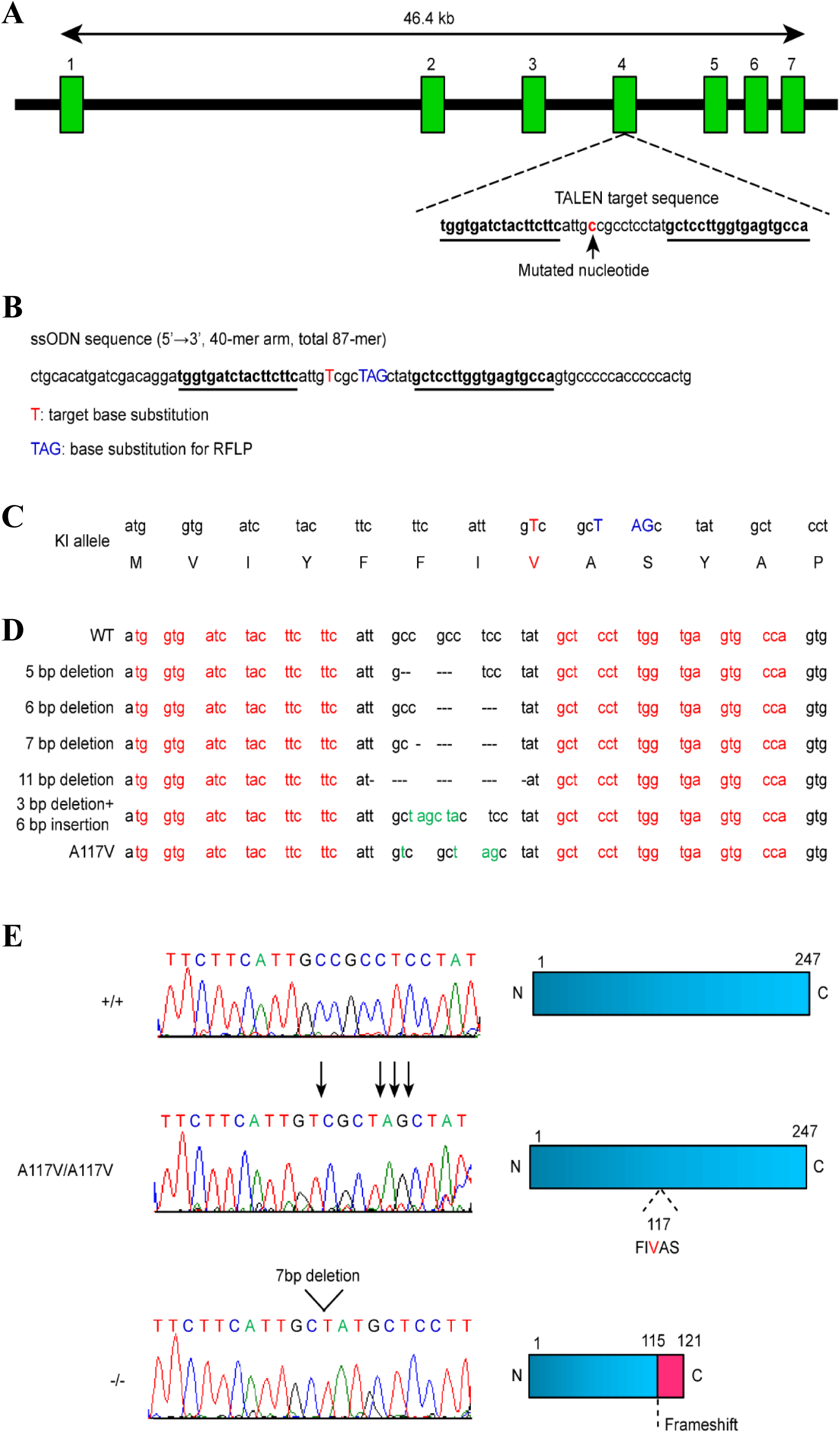
Generation of *Mmd2* A117V knock-in mice using Platinum TALEN. Panel A shows the genomic structure of *Mmd2*, indicating the two binding sites of the TALENs. TALEN pairs were designed to bind the exon 4 in the *Mmd2* gene. Panel B specifies the ssODN sequence. Panel C indicates the KI allele. Panel D shows the sequence information of *Mmd2* mutant alleles, specifically, sequences obtained from mutant mice, which were generated by microinjection of TALEN mRNA. The DNA sequences that were used for designing the TALENs are highlighted in red. Nucleotide mutations and indels are shown. Panel E illustrates the Sanger sequencing performed to confirm the *Mmd2* variant. The reference nucleotide C was substituted with variant nucleotide T in the mutant sample. A 7 bp deletion resulted in frameshifting and thus in truncated proteins.

### *Mmd2* mutation causes severe alveolar bone loss

Whether severe periodontitis was induced in *Mmd2*^A117V/A117V^ and *Mmd2*^-/-^ mice was investigated using the common ligature-induced periodontitis model (Abe, 2013). Alveolar bone was only markedly absorbed at the time of inflammation in both *Mmd2*^A117V/A117V^ and *Mmd2*^-/-^ mice, compared to wild-type (*Mmd2*^+/+^) mice (Fig. 5A). The degree of alveolar bone loss was statistically higher in *Mmd2*^A117V/A117V^, *Mmd2*^A117V/+^, and *Mmd2*^-/-^ mice, than in *Mmd2*^+/+^ mice (p = 0.01) (Fig. 5B).

**Fig. 5.**
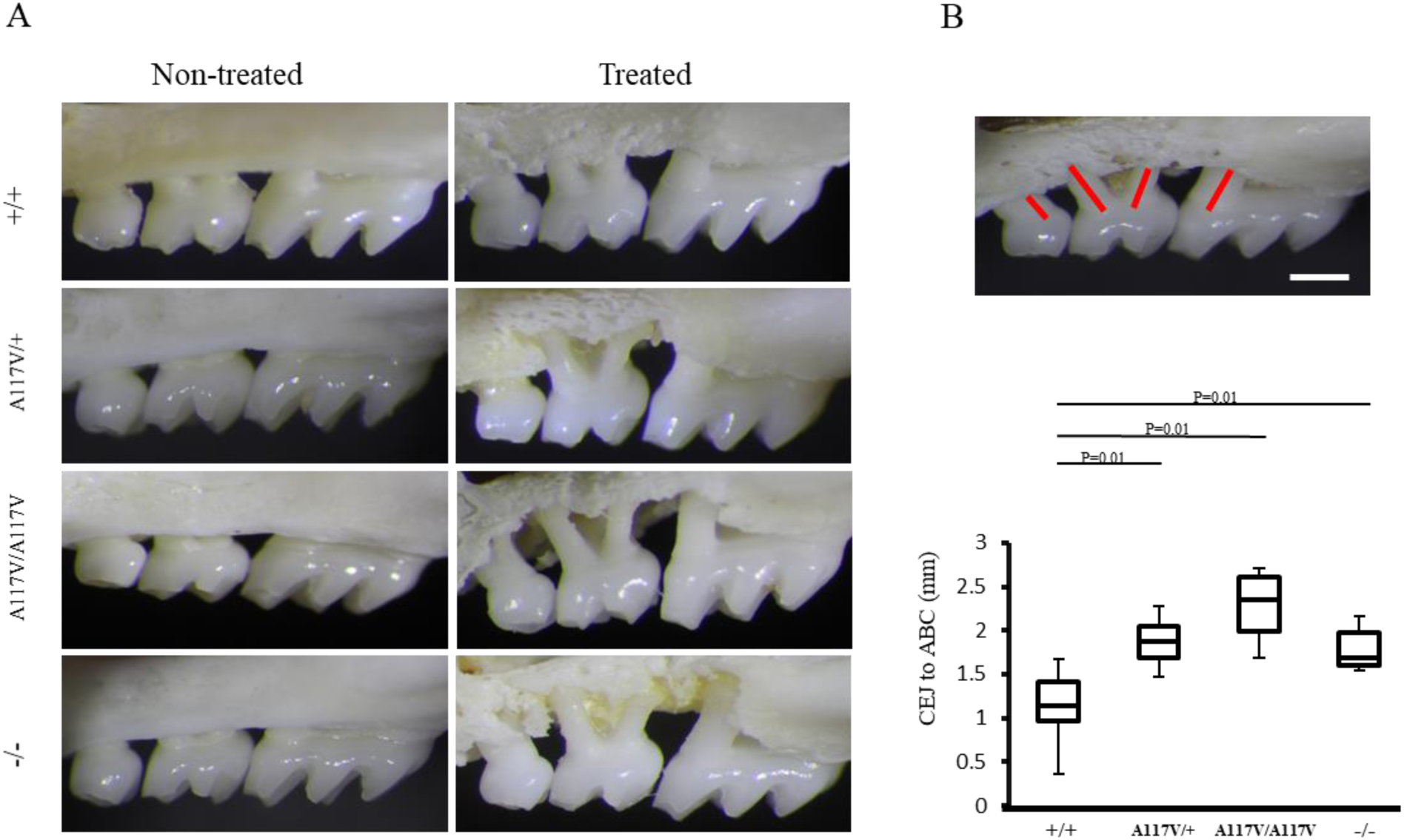
Periodontitis induction in *Mmd2*^A117V/A117V^ and *Mmd*2^-/-^ mice. Panel A shows representative photographs of the maxilla of treated and non-treated mice. Scale bar = 1 mm. Panel B displays schematics of periodontal bone loss measurements. The distance between cement-enamel junction and alveolar bone crest at the distal facial side of the first molar, at the mesial and distal facial side of the second molar, and at the mesial facial side of the third molar were measured (+/+, n = 19; A117V/+, n = 20; A117V/A117V, n = 8; -/-, n = 8). The results are expressed as the mean ± standard deviation. Those p-values ≤ 0.05 were considered statistically significant by using Student’s t test.

### *Mmd2* mutation causes abnormal differentiation of granulocytes

We also examined whether the differentiation of HSPCs to granulocytes was prevented in *Mmd2*^A117V/A117V^ and *Mmd2*^-/-^ mice. Blood parameters were analyzed to determine whether *Mmd2*^A117V/A117V^ and Mmd2^-/-^ mice showed mild neutropenia. The number of red blood cells and platelets, as well as the levels of hemoglobin did not change in *Mmd2*^A117V/A117V^ and *Mmd*2^-/-^ mice (Table S4). However, the ratios of granulocytes in these two mice were lower than that in *Mmd2*^+/+^ mice (*Mmd2*^A117V/A117V^ mice: p = 0.006 and *Mmd2*^-/-^ mice: p = 0.03) (Fig. 6A). To test whether this decrease in granulocyte numbers was associated with the differentiation ability of HSPCs, we examined the colony-forming ability of bone marrow cells in *Mmd2*^A117V/A117V^ and *Mmd2*^-/-^ mice. Stimulation with interleukin (IL)-3, granulocyte and macrophage colony stimulating factor (GM-CSF), and macrophage colony stimulating factor (M-CSF) indicated an abundance of early myeloid or monocytic precursor cells in *Mmd2*^A117V/A117V^ and *Mmd2*^-/-^ bone marrow cells. In contrast, stimulation of *Mmd2*^A117V/A117V^ and *Mmd2*^-/-^ bone marrow cells with G-CSF resulted in fewer colonies than those obtained with *Mmd2*^+/+^ bone marrow cells (p < 0.001) (Fig. 6B). This suggested that the number of granulocytic precursor cells decreased in the bone marrow of *Mmd2*^A117V/A117V^ and *Mmd2*^-/-^ mice. Moreover, neutrophil chemotaxis, assessed by stimulation with fMLP, also decreased in *Mmd2*^A117V/A117V^ and *Mmd2*^-/-^ mice compared with that in *Mmd2*^+/+^ mice (Fig. 6C).

**Fig. 6.**
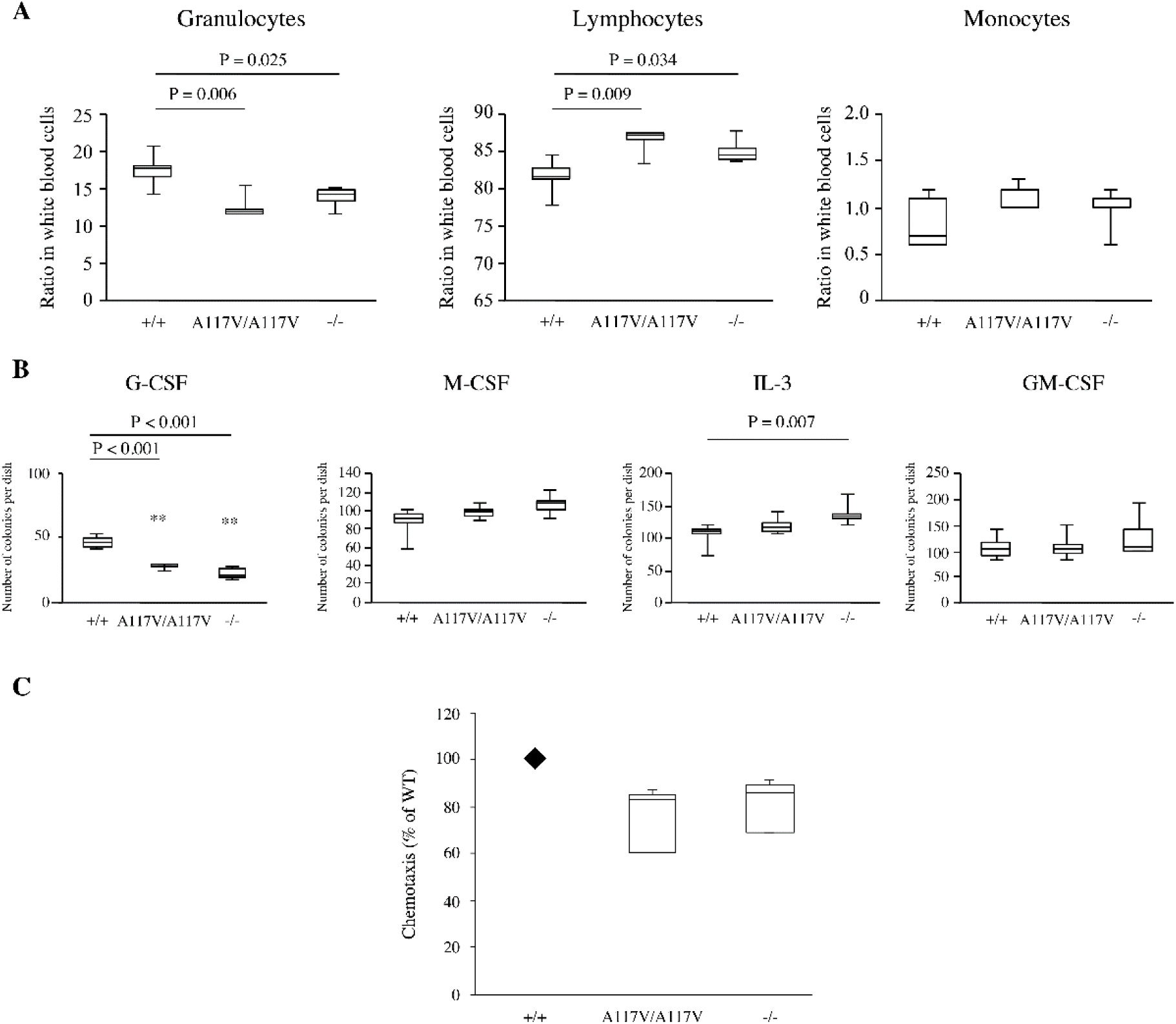
Cellular analysis of *Mmd2*^A117V/A117V^ and *Mmd*2^-/-^ mice. Panel A shows ratios of granulocytes, lymphocytes, and monocytes in white blood cells (n = 5). Panel B indicates colony formation from bone marrow cells in response to cytokines (n = 6). The results are the average of at least six independent experiments performed with each individual mouse. T bars indicate standard deviations. Panel C reveals that neutrophil chemotaxis, induced by fMLP (100 nM, 4 h), in *MMD2*^A117V/A117V^ and *MMD2*^-/-^ mice was decreased compared with that of *MMD2*^+/+^ mice (n = 5). The results are expressed as the mean ± standard deviation. Those p-values ≤ 0.05 were considered statistically significant by using Student’s t test.

Thus, severe alveolar bone loss and abnormal differentiation of granulocytes in both *Mmd2*^A117V/A117V^ and *Mmd2*^-/-^ mice strongly suggested that *MMD2* mutation in our patients could be the cause of aggressive periodontitis with neutropenia.

## Discussion

Study of subjects from a Japanese family indicated that *MMD2* gene was involved in autosomal dominant aggressive periodontitis. Our findings strongly suggested that *MMD2* gene was involved in the differentiation and function of neutrophils and that the presence of mutations in *MMD2* reduced the protective response to chronic bacterial infection. In patients with mutations in *MMD2* gene, HSPCs cannot differentiate into granulocytes and thus they may leak from bone marrow into peripheral blood. *MMD2* mutation was also associated with a decrease in neutrophil numbers and dysfunction, leading to severe periodontal destruction.

Neutrophil abnormalities are also found in severe congenital neutropenia (SCN). SCN, which poses a severe risk of infection since neonatal age, originates as a result of mutations in one of several different genes (Lanciotti et al., 2009; Lundén et al., 2009; Klein et al., 2007; Karsunky et al., 2002; Person et al., 2003; Boztug et al., 2009). These genes also play a role in the differentiation and function of neutrophils that are produced in the bone marrow. SCN has been shown to cause severe periodontal tissue destruction in addition to systemic infection, while aggressive periodontitis patients with mutations in *MMD2* are healthy but have localized infections in the oral cavity. This study showed that neutrophil abnormalities were important for the development of periodontal disease, because periodontal tissue was destroyed even if the neutropenia was mild. Additionally, aggressive periodontitis with *MMD2* mutation was considered to belong to the same spectrum of SCN due to a common mechanism that leads to abnormal number and function of neutrophils.

At present, symptomatic treatment is given for aggressive periodontitis; however, it does not fully recover the periodontal tissue. There are reports of SCN patients with absolute neutrophil counts normalized by G-CSF who still have severe periodontitis (Putsep, 2002; Carlsson et al., 2006). Therefore, the level of absolute neutrophil count normalized by G-CSF is not enough to maintain normal oral health in these patients. A more detailed examination of the immune response for aggressive periodontitis caused by *MMD2* mutation may lead to the development of a new treatment alternative to not only aggressive periodontitis caused by *MMD2* mutation but also to periodontitis in general.

MMD and MMD2 are two members of the progestin and adipoQ receptor family (Tang et al., 2005). As its name suggests, MMD is involved in macrophage activation and may increase the production of TNF-α and nitric oxide in lipopolysaccharide-stimulated macrophages through ERK1/2 and Akt phosphorylation (Liu et al., 2012). A genome-wide association study in patients with Crohn’s disease (CD) identified MMD2 as a CD-related gene (Montero-Melendez, 2013). CD is an inflammatory disease due to abnormal immune reaction to many commensal bacteria in genetically susceptible individuals. Thus, *MMD2* gene may be involved in the immune response system to chronic bacterial infection. Therefore, in presence of the *MMD2* mutation and harmful bacteria, diseases in the intestine and the oral cavity are likely to develop.

Additionally, identification of *MMD2* may allow to differentiate between chronic and aggressive periodontitis, as until today there is a controversy about whether chronic and aggressive periodontitis should be classified or not as the same type of periodontitis. As a conclusion, our study highlighted the influence of mild immune system defects on the onset of aggressive periodontitis, which should be considered during the diagnosis of the disease. Furthermore, the study of mild neutropenia and related diseases may attract the attention of the medical field other than periodontology and lead to the development of new diagnostic and therapeutic methods.

## Materials and Methods

### Study family

This study was approved by the Human Subjects Committees of Hiroshima University. Written informed consent was obtained from all subjects. All affected individuals were diagnosed with aggressive periodontitis according to periodontal and X-ray examinations. Blood was collected from the four affected and the four unaffected individuals in this family for genetic analyses. Also, blood and bone marrow samples were collected from III-2 and III-4 patients, and FACS analysis, chemotaxis assay, and CT imaging were performed.

### Differentiation of human CD34^+^ HSPCs into granulocytes

Cells were purified from the bone marrow and peripheral blood. Mononuclear cells, previously separated by Ficoll-Hypaque density centrifugation (GE Healthcare Biosciences, Uppsala, Sweden), were stained with a variety of antibodies and subjected to flow cytometry. CD34^+^ HSPCs were isolated from patients’ blood using the CD34 Microbeads Kit (Miltenyi Biotec, Bergisch Gladbach, Germany). CD34^+^ HSPCs from healthy volunteers were purchased from HemaCare (HemaCare, Northridge, CA, USA). Then, CD34^+^ HSPCs were incubated in StemSpan SFEM II medium with StemSpan Myeloid Expansion Supplement (Stem cell technologies, Vancouver, Canada) for 10 days. Further, harvested cells were stained with anti-CD33 antibody (Bio Biosciences, Franklin Lakes, NJ, USA) and subjected to flow cytometry analysis.

### Chemotaxis assay

Neutrophils were suspended in RPMI1640 (Nakalai tesque, Kyoto, Japan) with 1000 mg/L glucose and penicillin/streptomycin. Chemotaxis was induced with fMLP (100 nM) for 120 min at 37 °C and measured by the Boyden chamber method with a 96-well micro-chemotaxis chamber containing a 3-µm pore-sized filter (CELL BIOLABS, INC., San Diego, CA, USA).

### Linkage analysis

The samples used for linkage analysis were II-1, II-6, II-9, III-2, III-3, III-4, and III-5. Genomic DNA (gDNA) was extracted from the venous blood of each individual according to standard protocols. We used the Genome-Wide Human SNP Array 6.0 (Affymetrix, Santa Clara, CA, USA) for genotyping single nucleotide polymorphisms (SNPs), and linkage analysis was performed by Allegro software, assuming dominant inheritance.

### Exome sequencing and variant filtering

The gDNA libraries were prepared using a SeqCap EZ Human Exome Library v2.0 (Roche, Basel, Switzerland). Sequencing was performed with 100-bp paired-end reads by the HiSeq2000 sequencer (Illumina, San Diego, CA, USA). We used Burrows-Wheeler Aligner for alignment and mapping, and SAMtools and Picard for SAM/BAM. Exome sequencing was performed using the GATK and SAMtools for variant calls and Annovar for annotation. Functional predictions of amino acid changes were performed using PolyPhen-2, Mutation Taster, SIFT, and the Combined Annotation Dependent Depletion (CADD). Control exome sequences were obtained from Japanese patients undergoing exome sequencing analysis for other diseases. All reported genomic coordinates were in GRCh37/hg19. The identified mutation was confirmed by standard polymerase chain reaction-based amplification, followed by sequence analysis using Applied Biosystems 3130 DNA sequencer (Thermo Fisher Scientific, Waltham, MA, USA).

### Double-staining procedures for immunofluorescence

The primary antibodies used in this study were anti-MMD2 (CUSABIO, Houston, TX) and anti-golgin-97 (Gene Tex, Irvine, CA, USA). Neutrophil-like HL-60 cells (1.0 × 10^5^ cells/well in chamber slides) were fixed in 4 % paraformaldehyde and permeabilized with 0.2 % Triton X-100. For immunofluorescence analysis, cells were assessed on cytospin preparations. Slides were incubated with anti-MMD2 and golgin-97 antibodies at 4 °C overnight. MMD2 and golgin-97 proteins were detected after incubation with Alexa Fluor-594 rabbit anti-donkey IgG and Alexa Fluor-488 mouse anti-donkey IgG secondary antibodies, respectively. Nuclei were stained with 4,6-diamidino-2-phenylindole (DAPI). Fluorescence signals were detected with Olympus Fluoview FV1000 laser scanning confocal microscope (Olympus, Tokyo, Japan).

### Generation of mouse model using Platinum TALEN gene editing

Mice carrying the *Mmd2* A117V variant were generated using the TALEN gene editing tool. The TALEN pair that showed a high targeting efficiency and low off-target effects was used for *in vitro* transcription by using the MEGAscript T7 Transcription Kit (Thermo Fisher Scientific, Yokohama, JAPAN). TALEN mRNAs were combined with the ssODN construct and injected into pronuclei of C57BL/6 single cell mouse embryos. We then backcrossed *Mmd2*^A117V/A117V^ and *Mmd*2^-/-^ mice with C57BL/6 mice for eight generations.

### Ligature-induced periodontitis

Periodontal inflammation and bone loss in a ligature-induced periodontitis model was initiated by the abundant local accumulation of bacteria on ligated molar teeth. To this end, a 5-0 silk ligature was tied around the maxillary second molar in 8-week-old male mice. The distance between cement-enamel junction and alveolar bone crest was examined at 7 days after placement of the ligatures.

### Colony formation assays

Mouse bone marrow cells (2.5×10^4^) suspended in methylcellulose semisolid medium (Methocult M3231) (Stem Cell Technologies) were plated in 35-mm culture dishes in the presence of 0.5 % FBS, 10 ng/mL mouse G-CSF, 10 ng/mL mouse GM-CSF, 10 ng/mL mouse M-CSF, and 10 ng/mL mouse IL-3 (BioLegend, San Diego, CA, USA).

### Statistical analysis

The results are expressed as the mean ± standard deviation. Statistical differences between the mean values of the control and experimental groups were analyzed by using Student’s t test. Those p-values ≤ 0.05 were considered statistically significant.

## Acknowledgments

We thank the families involved in this research. We would like to thank Editage (www.editage.jp) for English language editing.

## Author contributions

Noriyoshi Mizuno, Hiroyuki Morino, and Keichiro Mihara: study concept and design, acquisition, analysis, and interpretation of data, manuscript preparation and revision; Tomoyuki Iwata: statistical analysis; Yoshinori Ohno, Shinji Matsuda, Kazuhisa Ouhara, Mikihito Kajiya, Kyoko Suzuki-Takedachi: data acquisition and patient evaluation; Yusuke Sotomaru, Katsuhiro Takeda, Shinya Sasaki, and Ai Okanobu: data acquisition and manuscript revision; Tetsushi Sakuma and Takashi Yamamoto: data acquisition; Yukiko Matsuda, Ryousuke Ohsawa, and Tsuyoshi Fujita: data analysis; Hideki Shiba, Hideshi Kawakami, Hidemi Kurihara: study concept and design, and manuscript revision.

## Additional Files

**Figure 2-source data 1.** Source data for Figure 2A.

**Figure 2-source data 2.** Source data for Figure 2C.

**Figure 5-source data 1.** Source data for Figure 5B.

**Figure 6-source data 1.** Source data for Figure 6A.

**Figure 6-source data 2.** Source data for Figure 6B.

**Figure 6-source data 3.** Source data for Figure 6C.

**Table 4 - source data 1.** Source data for Table 4.

## Tables

**Table S1.** Data of complete blood count (Patients).

| **Table S1. Data of complete blood count (Patients).** |  |  |  |  |
| --- | --- | --- | --- | --- |
|  | III-2 | III-3 | III-4 | Reference Range |
| RBC (/nl) | 4,960 | 4,870 | 4,840 | 4,200-5,600 |
| Hemoglobin (g/dl) | 15.3 | 14.6 | 14.1 | 12.5-17.0 |
| PLT (nl) | 202 | 248 | 208 | 150-350 |
| WBC (/nl) | 3,120 | 2,500 | 3,080 | 4,000-9,000 |
| Neutrophil (%) | 31.1 | 35.3 | 31.3 | 43.0-71.0 |
| Lymphocyte (%) | 55.9 | 53.7 | 57.6 | 30.0-41.0 |
| Monocyte (%) | 5.8 | 8.0 | 6.6 | 3.0-6.0 |
| Eosinophil (%) | 5.9 | 2.0 | 3.2 | 2.0-6.0 |
| Basophil (%) | 1.3 | 1.0 | 1.3 | 0.0-2.0 |
| Neutrophil (μl) | 970 | 883 | 964 | 1,720-6,390 |

**Table S2.** Summary of exome sequencing.

| **Table S2. Summary of exome sequencing.** |  |  |  |
| --- | --- | --- | --- |
| Sample | III-2 | III-3 | III-4 |
| <Sequence Info> |  |  |  |
| Total reads (bp) | 7,835,611,079 | 7,647,632,797 | 7,246,927,551 |
| Mean depth | 73.9 | 72.1 | 68.3 |
| >10 reads (%) | 61.7 | 61.3 | 61.5 |
| <Variant> |  |  |  |
| Total | 91,643 | 90,396 | 91,052 |
| -dbSNP/TGP | 13,818 | 13,516 | 13,128 |
| Exon/splice | 2,148 | 2,122 | 1,958 |
| AA change | 1,743 | 1,725 | 1,589 |
| Het | 1,547 | 1,534 | 1,392 |
| Common |  | 282 |  |
| Disease specific |  | 7 |  |
| Pathogenic |  | 3 |  |
| Linkage |  | 1 |  |

**Table S3.** Candidate variants.

| Table S3. Candidate variants. |  |  |  |  |  |  |  |
| --- | --- | --- | --- | --- | --- | --- | --- |
| Variant | Gene | PolyPhen-2 | Mutation Taster | SIFT | CADD | Segregation | Linkage |
| Chr1:236144983C>T | *NID1* | PROBABLY DAMAGING | DISEASE CAUSING | DAMAGING | 27.7 | No | No |
| Chr3:195506098T>G | *MUC4* | BENIGN | POLYMORPHISM | NOT SCORED | 5.977 | No | No |
| Chr3:195508545->G | *MUC4* | N.A. | DISEASE CAUSING | DAMAGING | 24.3 | No | No |
| Chr6:30954525G>A | *MUC21* | BENIGN | POLYMORPHISM | TOLERATED | 0.007 | No | No |
| Chr6:30954526C>T | *MUC21* | BENIGN | POLYMORPHISM | TOLERATED | 5.61 | No | No |
| Chr7:4955653G>A | *MMD2* | PROBABLY DAMAGING | DISEASE CAUSING | DAMAGING | 28.9 | Yes | Yes |
| Chr12:90908G>A | *LOC100288778* | BENIGN | N.A. | TOLERATED | 14.68 | No | No |
| N.A.: not available |  |  |  |  |  |  |  |

**Table S4.** Data of complete blood count (mouse).

| **Table S4. Data of complete blood count (mouse).** |  |  |  |
| --- | --- | --- | --- |
|  | MMD2<sup>+/+</sup> | MMD2<sup>A117V/A117V</sup> | MMD2<sup>-/-</sup> |
| RBC ($\times 10^4/\mu\text{l}$) | 908.0 ± 24.8 | 884.8 ± 99.5 | 901.3 ± 48.0 |
| Hemoglobin (g/dl) | 14.0 ± 0.9 | 14.6 ± 1.8 | 14.0 ± 0.9 |
| PLT (/nl) | 78.9 ± 19.1 | 100.4 ± 46.0 | 86.5 ± 26.7 |
| WBC (/μl) | 2398 ± 999 | 1738 ± 316 | 1598 ± 361 |

The results are expressed as the mean ± standard deviation.

